# Exploring the Interdependence of TKS4 and CD2AP: Implications for EMT Process and Early Detection in Colon Cancer

**DOI:** 10.1101/2023.01.13.523903

**Authors:** Anita Kurilla, Loretta László, Tamás Takács, Álmos Tilajka, Laura Lukács, Julianna Novák, Rita Pancsa, László Buday, Virag Vas

**Author notes:** Correspondence: Anita Kurilla Virag Vas.

## Abstract

**Background:** Colon cancer is a leading cause of death worldwide. Although several biomarkers have been developed, more sensitive and specific methods are needed for its early detection. TKS4 and CD2AP scaffold proteins have been already linked to dynamic actin assembly-related processes, such as colon cancer cell migration, although their co-instructive role during tumour-formation remains unknown. Therefore, this study was designed to validate TKS4-CD2AP interaction and study the interdependent effect of TKS4/CD2AP on oncogenic events.

**Methods:** CD2AP was identified as a novel TKS4 interacting partner in six cell lines via co-immunoprecipitation-mass spectrometry. The interaction was validated via western blot, immunocytochemistry, DuoLink assay and peptide microarray. Gene silencing and overexpressing experiments were performed to uncover the cooperative effects of TKS4 and CD2AP in cell movement and in epithelial-mesenchymal transition (EMT). The expression levels of TKS4 and CD2AP mRNAs were quantified using a human colon cancer array, and the results were subjected to bioinformatic database analysis.

**Results:** The molecular biology analysis revealed that one of the SH3-domain of CD2AP interacts with a proline-rich short linear motif in TKS4. Functional studies showed that TKS4 and CD2AP form a scaffold complex that regulates migration- and EMT-related pathways of colon cancer cells. The relative TKS4 and CD2AP expression level measurements pointed out that CD2AP/TKS4 ratio is a sensitive biomarker for the identification of tumorous colon tissue.

**Conclusions:** This is the first study demonstrating the TKS4-CD2AP protein-protein interaction in vitro and in vivo and their interdependent regulative effect on mesenchymal transition-like process in colon cancer. Furthermore, the results highlight that the relative expression levels of CD2AP and TKS4 might serve as a biomarker for the diagnosis of early-stage colon cancer.

## 1. Introduction

Colon (colorectal) cancer is one of the most common cancer types and is the second leading cause of cancer death worldwide. Mortality and survival rates mostly depend on appropriate screening and treatment techniques. As these methods have improved in the last decade, death rates have dropped significantly (seer.cancer.gov). Currently, several blood-based and molecular biomarkers have been developed and used clinically, including the presence of *KRAS* mutations as well as the presence of circulating tumour DNA (ctDNA) or circulating tumour cells (CTCs). However, more sensitive and specific methods are needed to detect colon cancer at its early stages (Jelski and Mroczko 2020).

Several molecular events are known to drive colon cancer formation, including the accumulation of oncogenic mutations, development of drug resistance, and initiation of the cellular invasion cascade, leading to the appearance of CTCs and metastasis (László et al. 2021). However, most of the molecular mechanisms underlying oncogenesis are not yet fully understood; therefore, identifying novel molecular players that regulate colon cancer invasiveness will help us understand its formation and progression, ultimately contributing to the discovery of novel biomarkers and therapeutic targets. The EMT process is the central regulator of metastasis during which epithelial cells transform into the mesenchymal state (Ye and Weinberg 2015). Although EMT has been generally considered as a binary process, lately it has been proven that rather a gradual transformation event takes place during transition. In this spectrum, EMT includes an intermediate partial EMT (pEMT) state in which cells maintain both epithelial and mesenchymal phenotypes, implying that co-expression of epithelial and mesenchymal markers can be observed (Yang et al. 2020). It is known that tumour cells can be heterogenous with epithelial, mesenchymal, and hybrid E/M (epithelial/mesenchymal) phenotypes simultaneously. Likewise, tumours with pEMT phenotype can also show intra-tumoral heterogeneity and epithelial-mesenchymal plasticity. The pEMT process includes different steps, such as early, intermediate, and late states that cover the spectrum from epithelial to completely mesenchymal phenotype (Pastushenko et al. 2018). Furthermore, pEMT has a high clinical significance as it was associated with worse prognosis and higher metastatic risk (Saitoh 2018) (Yagasaki et al. 1996). Presently, there is no clear consensus in defining pEMT phenotype, however, co-expression of different epithelial and mesenchymal markers is considered as an intermediate cell state (Aggarwal et al. 2021).

Recently, CD2-associated protein (CD2AP) and tyrosine kinase substrate with four SH3 domains (TKS4) have emerged as potential cancer formation-regulating factors. These two proteins, which lack enzymatic activity, are scaffolds/adaptors that serve as molecular hubs within cells by binding and releasing multiple signaling molecules via phosphorylation-dependent or -independent interactions mainly mediated by SH3 domains and proline-rich regions (Figure 1a). Although their roles in an EMT-like process have been studied separately, an interaction or co-regulation involving TKS4 and CD2A has not been reported. We suspected the possibility of interplay based on a previous study (Bazalii et al. 2015), which demonstrated that TKS4 interacts with a close paralog of CD2AP called CIN85 (SH3KBP1). However, this study only suggested that the interaction is mediated by an SH3 domain in CIN85, but did not identify the exact interacting residues within TKS4. CD2AP and CIN85 each have three SH3 domains, which are special in the sense that they recognize non-canonical PxP/A/V/IxPR motifs instead of the canonical PxxP SH3 binding sites (Kowanetz et al. 2003)(Rouka et al. 2015) (Figure 1a). CD2AP was originally implicated in dynamic actin remodeling via direct binding to either filamentous actin (Lehtonen et al. 2002) or cortactin, an actin-binding scaffold protein (Lynch et al. 2003). Later, it was found that CD2AP also plays a regulatory role in epithelial cell junction formation (Tang and Brieher 2013), and it was also linked to gastric cancer development as it can inhibit metastasis by promoting cellular adhesion and cytoskeleton assembly (Xie et al. 2020).

**Figure 1.**
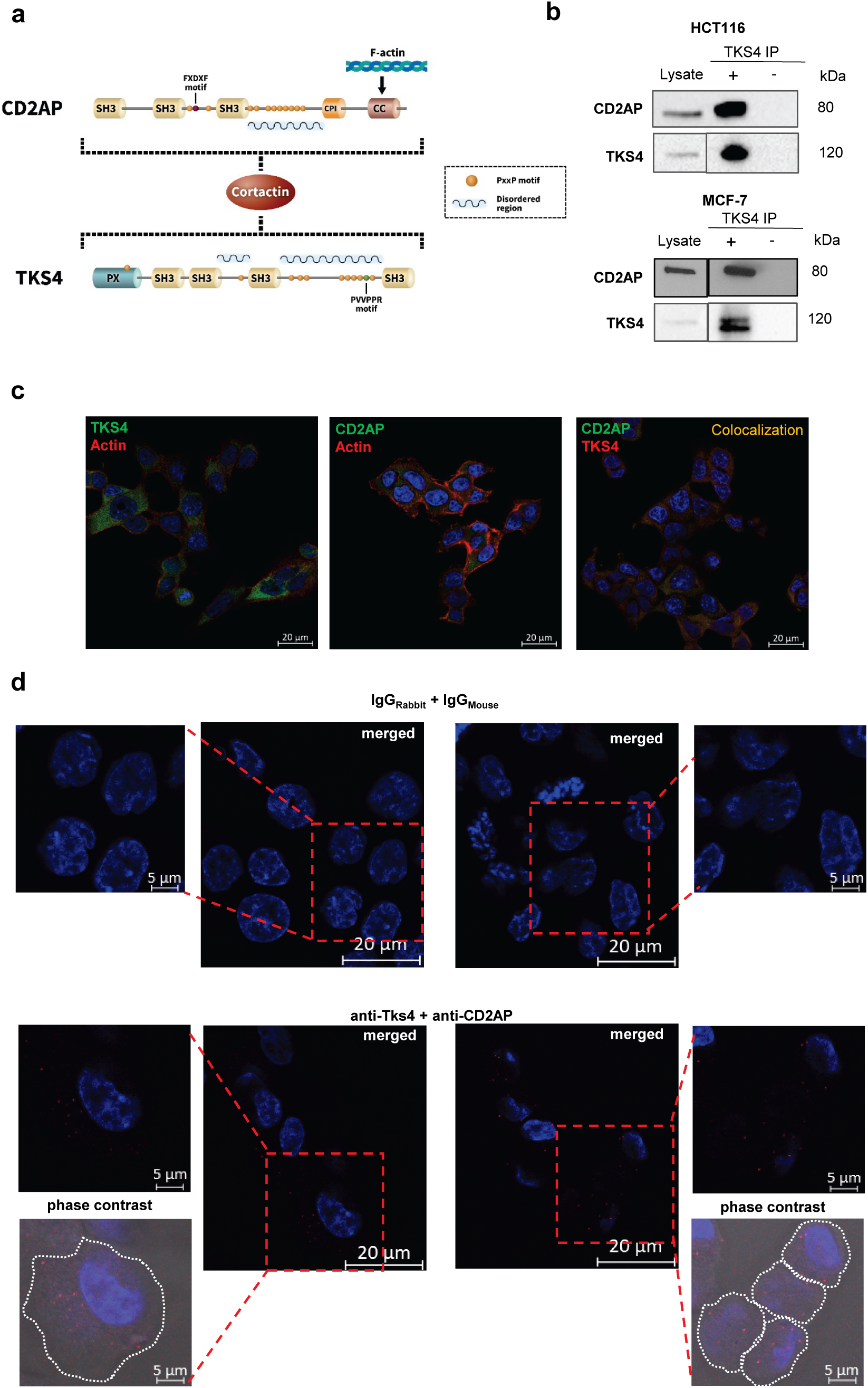
A combination of co-IP and MS was used to identify CD2AP and TKS4 interaction, and the validation of CD2AP-TKS4 interaction via IP-WB, DuoLink Proximity Ligation Assay and Peptide array. **(a)** Schematic representation of CD2AP and TKS4 proteins highlighting the potential (proline rich) interacting motifs in the molecules (FxDxF, PXXP, PVVPPR) and the common binding partner cortactin. Domain names in the proteins are the following: CPI: capping protein interaction motifs, CC: coiled-coil region, SH3: SRC homology 3 domain, PX: phosphoinositide-binding domain. **(b)** Validation of CD2AP as a TKS4-interacting partner via IP-WB in HCT116 and MCF-7 cells. Labeling is the following: *Lysates*: whole cell lysates isolated from HCT116 and MCF-7 cells; *TKS4 IP+*: immunoprecipitated protein lysates with TKS4-specific antibody; *TKS4 IP−*: immunoprecipitation-control without anti-TKS4 antibody. **(c)** Immunocytochemistry of TKS4 and CD2AP single staining (Alexa 488-green, and counterstained Phalloidin-actin) and double staining for CD2AP (Alexa 488 (green)) and for TKS4 (Alexa 546 (red)) of HCT116 cells. The colocalization analysis of the two proteins showed overlap as orange-fluorescent signals. **(d)** Detecting the CD2AP-TKS4 interaction in HCT116 cells via DuoLink assay counterstained with DAPI. In control samples (stained with IgG_Rabbit_ and IgG_Mouse_ antibodies) no fluorescence signal was generated. In anti-TKS4 and anti-CD2AP antibodies-stained samples, the two DPLA probes are in proximity resulted in as red-fluorescent dots. Colocalization was present in the cytoplasm and under the cell membrane seen on merged phase contrast images.

TKS4 also serves as a docking site for molecules involved in cell movement and actin rearrangement, e.g., cortactin or N-WASP, and it regulates podosome formation in normal cells and invadopodia formation in cancer cells (Buschman et al. 2009)(Iizuka et al. 2016) (Figure 1a). The involvement of TKS4 in EGF-EGFR-SRC signaling has been previously described, and regulatory roles in mesenchymal stem cell differentiation, adipose tissue, and bone homeostasis have recently been discovered (Dülk et al. 2016)(Vas et al. 2019a)(Vas et al. 2019b). Recent progress in TKS4-related research has revealed that TKS4 might also be implicated in several cancer types, including breast cancer (Maziveyi et al. 2018), melanoma (Iizuka et al. 2016), acute myeloid leukemia (Li et al. 2019), and prostate cancer (Caires-dos-Santos et al. 2020). Moreover, it was recently demonstrated in a colon cancer cell line that reduced TKS4 level leads to disturbed EMT-like processes (Szeder et al. 2019).

Based on the notion that CD2AP can also bind TKS4 and the fact that both proteins have been implicated in cancer cell biology and metastatic process raises the possibility that their interaction is fundamental to tumour development. Here, we validated that CD2AP and TKS4 can form a complex in several different cancer cell lines. Furthermore, we applied functional assays to test the interdependent effect of TKS4/CD2AP on oncogenic events. SiRNA-based silencing was used to downregulate their transcript levels and overexpressing experiments were performed in HCT116 colon cancer cells to measure the resulting changes in EMT process-related gene expression levels and cell migration. The individual and relative expression levels of CD2AP and TKS4 have been also analysed in human colon cancer samples in a tissue array and in databases to shed light on the potential effects of their interplay during colon cancer formation.

We found that CD2AP and TKS4 individually and interdependently inhibit pEMT and impair colon cancer cell migration. We found that the ratio of CD2AP and TKS4 expression levels has an instructive role in tumour phenotypes and that CD2AP/TKS4 ratios are outstanding biomarkers for the identification of tumorous colon tissue.

## 2. Materials and Methods

### 2.1. Cell culture

All cell lines were obtained from ATCC and were tested for mycoplasma contamination using MycoAlert Mycoplasma Detection Kit (Lonza, Basel, Switzerland). HCT116 (RRID:CVCL_0291) cells were grown in McCoy’s 5A Medium (Thermo Fisher Scientific, 16600082). DLD-1 (RRID:CVCL_0248) cells were grown in RPMI medium. MCF-7 (RRID:CVCL_0031), HEK293 (RRID:CVCL_0045), HPAC (RRID:CVCL_3517), and NIH3T3 (RRID:CVCL_0594) cells were grown in DMEM. All media were supplemented with 10% FBS (Thermo Fisher Scientific, 26400044) and penicillin/streptomycin, and cells were incubated at 37 °C in 5% CO_2_.

### 2.2. siRNA and DNA plasmid transfection

Small interfering RNA (siRNA) transfection was performed with Lipofectamine RNAiMax Transfection Reagent (Thermo Fisher Scientific, 13778030) according to the manufacturer’s instructions. SiRNAs targeting CD2AP (133808) and TKS4 (252894) were purchased from Thermo Fisher Scientific and used at 10 nM. Scrambled siRNA was used as a negative control at 10 nM. After transfection, cells were incubated for 48 h prior to RNA or protein isolation.

For DNA plasmid transfection, V5-TKS4 or/and MYC-CD2AP (Myc-DDK-Tagged CD2AP clone) (CAT#: RC210191, Origene Technologies Inc., Rockville, MD) plasmids were transfected using X-tremeGENE Hp DNA Transfection Reagent (Roche) following the manufacturer’s instructions. V5-TKS4 plasmid contained the coding sequence of human TKS4 in the pcDNA3.1/TOPO-V5-His vector (Lányi et al. 2011). During experiments, negative control wells were treated with transfection reagent only. After transfection, cells were incubated with transfection reagent for 48 h before cell lysis. All experiments were repeated three times.

### 2.3. RNA isolation and quantitative real-time polymerase chain reaction (qRT-PCR)

RNA was extracted using Direct-zol RNA Miniprep Kit (Zymo Research). First Strand cDNA Synthesis Kit (Roche) and Fast SYBR Green Master Mix (Thermo Fisher Scientific) were used in qRT-PCR (QuantStudioPro6).

Data were normalized to the *GAPDH* and *PUM1* transcript levels, and the 2^-ΔΔCt^ method was used to calculate relative gene expression levels. Primers used in this study are listed in Supplementary Table 2.

### 2.4. Western blotting and immunoprecipitation

For protein extraction, cells were first washed with ice-cold phosphate-buffered saline (PBS). Next, ice-cold harvest buffer (30 mM Tris pH 7.5, with 100 mM NaCl, 1% Triton X-100, 10 mM NaF, 1 mM Na_3_VO_4_, 1 mM EGTA, 2 mM 4-nitrophenyl phosphate, 10 mM benzamidine, 1 mM phenylmethylsulphonyl fluoride, 25 μg/mL Pepstatin A, 25 μg/mL trypsin inhibitor, and 25 μg/mL aprotinin) was added to the cells. Lysates were centrifuged at 14000 rcf, 10 min at 4 °C.

Immunoprecipitation was performed using Protein-A Sepharose from *Staphylococcus aureus* (Merck Millipore) and anti-TKS4 antibody (Lányi et al. 2011). 1.5 ml cell lysates, 40 µl agarose beads, and 10 µl polyclonal anti-TKS4 antibody were incubated for 1 hour at 4 °C and then washed three times with 1 ml PBS+0.4% Triton-X at 14000 rcf for 1 min at 4 °C.

Sample loading buffer (4.2 ml H20, 1 ml 6.8 pH Tris, 2.8 ml Glicerin, 1.6 ml 10% SDS, 0.4 ml β-mercaptoethanol, 1 mg bromophenol blue) was added to the supernatants, followed by incubation at 95 °C for 5 min. Each protein sample was subjected to 10% SDS-PAGE gel (BioRad) and blotted. Nitrocellulose membranes were blocked and incubated for 60 min with anti-TKS4 (Lányi et al. 2011),1:5000) and anti-CD2AP (clone 2A2.1, MABT419, 1:2000) antibodies at room temperature or overnight at 4 °C with anti-E-cadherin (ab15148 1:500), or anti-vimentin (ab137321 1:1000) antibody. After three washes, the membranes were incubated for 60 min with horseradish peroxidase-conjugated secondary antibody (GE Healthcare) and then washed four times for 15 min each. The proteins of interest were visualized using enhanced chemiluminescence (ECL) detection reagents (Amersham, Little Chalfont, UK). Chemiluminescent imaging was performed with a ChemiDoc MP system (Bio-Rad). The analysed protein levels were normalized to total protein staining.

### 2.5. Mass spectrometry analyses

After immunoprecipitation, samples were loaded onto 10% SDS-PAGE gels. Next, proteins were visualized via Coomassie Brilliant Blue dye, and gels were fractionated horizontally based on molecular weight (300-100 kDa, 100-50 kDa, 50-25 kDa, 25-10 kDa). Mass spectrometry analyses were performed by UD-GenoMed Medical Genomic Technologies Kft (Debrecen, Hungary).

#### In-gel digestion of proteins

The protein bands were excised from the gel and subjected to in-gel trypsin digestion. The bands were destained using a 1:1 ratio of 25 mM ammonium bicarbonate (pH 8.5) and 50% acetonitrile followed by the reduction of the proteins using 20 mM dithiothreitol (Sigma, St. Louis, MO, USA) for one hour at 56°C. The samples were further alkylated with 55 mM iodoacetamide (Sigma, St. Louis, MO, USA) for 45 minutes in dark. Overnight trypsin digestion was carried out with 100 ng stabilized MS grade trypsin (ABSciex, Framingham, MA, USA) at 37°C. The reaction was stopped by the addition of concentrated formic acid. The tryptic peptides were extracted from the gel pieces, dried in a vacuum concentrator (Thermo Scientific, Waltham, MA, USA) and kept at -20°C until the mass spectrometry analysis.

#### Liquid chromatography-mass spectrometry analysis

For protein identification by liquid chromatography with tandem mass spectrometry, the peptides were re-dissolved in 10 μl 1% formic acid (VWR Ltd., Radnor, PA, USA) and were separated in a 180 min water/acetonitrile gradient using an Easy nLC 1200 nano UPLC (Thermo Scientific, Waltham, MA, USA). The peptide mixtures were desalted in an ACQUITY UPLC Symmetry C18 trap column (20 mm × 180 µm, 5 μm particle size, 100 Å pore size; Waters, Milford, MA, USA), followed by separation in a nanoACQUITY Peptide BEH C18 analytical column (150 mm × 75 μm, 1.7 μm particle size, 130 Å pore size; Waters, Milford, MA, USA). The chromatographic separation was performed by using a gradient of 5–7% solvent B over 5 min, followed by a rise to 15% of solvent B over 50 min, and then to 35% solvent B over 60 min. Thereafter, solvent B was increased to 40% over 28 min and then to 85% over 5 min, followed by a 10 min rise to 85% of solvent B, after which the system returned to 5% solvent B in 1 min for a 16 min hold-on. Solvent A was 0.1% formic acid in LC water (Sigma, St. Louis, MO, USA); solvent B was 95% acetonitrile (Sigma, St. Louis, MO, USA) containing 0.1% formic acid. The flow rate was set to 300 nL/min.

Data-dependent acquisition experiments were carried out on an Orbitrap Fusion mass spectrometer (Thermo Scientific, Waltham, MA, USA). The 14 most abundant multiply charged positive ions were selected from each survey MS scan using a scan range of 350–1600 m/z for MS/MS analyses (Orbitrap analyzer resolution: 60.000, AGC target: 4.0e5, acquired in profile mode). Collision-induced dissociation (CID) fragmentation was performed in the linear ion trap with 35% normalized collision energy (AGC target: 2.0e3, acquired in centroid mode). Dynamic exclusion was enabled during the cycles (exclusion time: 45 s).

#### Protein identification

The acquired LC-MS/MS data were used for protein identification with the help of MaxQuant 2.0.1 software (Cox and Mann 2008) searching against the Human SwissProt database (release: 2020.06, 20394 sequence entries) and against the contaminants database provided by the MaxQuant software. Cys carbamidomethylation, Met oxidation and N-terminal acetylation were set as variable modifications. A maximum of two missed cleavage sites were allowed. Results were imported into Scaffold 5.0.1 software (ProteomeSoftware Inc., Portland, OR, USA). Proteins were accepted with at least 2 identified peptides using a 1% protein false discovery rate (FDR) and 95% peptide probability thresholds.

### 2.6. Wound healing assays

HCT116 cells were forward-transfected with siRNAs or plasmid DNA. Next, 5×10^5^ cells were cultured in Ibidi Culture-Insert well (35 mm, Cat.No:81176). After 24 hours, cells reached confluency and the silicone inserts were removed to induce uniform scratch in the cell layer. The dishes were filled with fresh McCoy’s Medium, and cells were maintained for 24 hours to close the wound. Images were acquired by Leica DMi1 Inverted Microscopy and using images from each triplicate wells the gap area was calculated. Experiments were repeated three times.

### 2.7. Immunocytochemistry

Cells were fixed with 4% PFA (Thermo Fisher Scientific) for 15 min at RT and then permeabilized with 0,1% TritonX-100 in sterile PBS for 10 min at RT, after blocking the samples with 5% BSA and 5% FBS in sterile PBS for 1 hour at RT. Next, cells were incubated with the appropriate primary antibody overnight at 4°C, and after several washing steps, secondary antibodies were used. The cell nuclei were stained with DAPI (Thermo Fisher Scientific). A Zeiss LSM-710 confocal microscopy system (Carl Zeiss Microscopy GmbH, Jena, Germany) was used to detect proteins of interest with a 63× objective. Images were analysed with ZEN 3.2 (Carl Zeiss Microscopy GmbH, Jena, Germany) and Image J software. Primary antibodies were the same as we used for western blot. Secondary antibodies were: Goat-Anti-Mouse antibody, Alexa Fluor 488, Thermo Fisher Scientific Cat# A-11029/, Goat anti-Rabbit IgG, Alexa Fluor 488 Thermo Fisher Scientific Cat # A-11008, Goat-Anti-Rabbit antibody, Alexa Fluor 546, Thermo Fisher Scientific Cat#A11035 for ICC analysis. Actin filaments were stained with CF®543 Phalloidin (Biotium, Cat: 00043). The colocalization coefficient was calculated with Image J-Ezcolocalization plugin according to this article (Stauffer et al. 2018).

### 2.8. DuoLink Proximity Ligation Assays

HCT116 cells were plated in 12-well Ibidi chambers and cultured for 2 days. Next, the cells were fixed in 4% PFA-PBS and further processed according to the manufacturer’s instructions (DUO92101−1KT, Sigma-Aldrich): permeabilization and blocking. The DPLA probe anti-rabbit plus binds to the TKS4 primary antibody (HPA036471, Sigma Aldrich, 1:100 dilution), whereas the DPLA probe anti-mouse minus binds to the CD2AP antibody (clone 2A2.1, Millipore, 1:250 dilution). The DPLA secondary antibodies generate a signal only when the two DPLA probes interact. Signal detection was conducted by ligation and rolling circle amplification with fluorescently labeled nucleotides. The amplified fluorescent DNA resulted in bright red dots, with each dot representing an individual CD2AP/TKS4 interaction event. Control samples were processed similarly without the addition of the primary antibodies. Staining and image acquisition were performed as previously described (Dülk et al. 2018). Briefly, nuclei were visualized via DAPI staining (with DuoLink mounting medium), and images were acquired on a Zeiss LSM710 inverted confocal microscope with a 40× objective (Carl Zeiss).

### 2.9. Peptide array

PepStar Peptide Microarray experiments were performed based on the method of Harnos and colleagues (2018) with some modifications (Harnoš et al. 2018). Microarrays were purchased from JPT (JPT Peptide Technologies GmbH). Peptides (15-mers with 11 overlapping residues) covering the complete TKS4 protein sequence (A1X283) were printed on a glass slide (25×75 mm). The peptide microarrays were printed in three identical subarrays. Several controls were also printed on the glass slides to exclude non-specific binding events or to confirm positive signals (Myc-tag, anti-Myc antibody, human serum albumin, human IgG, rabbit IgG). When screening the binding ability of CD2AP to TKS4 peptides, Myc-tagged CD2AP protein was used. Myc-DDK tagged human CD2-associated protein (RC210191) was purchased from Origene Technologies. The screening experiments were performed using two glass slides. As a control, one slide was incubated only with Alexa Fluor^®^ 488-conjugated Myc-Tag (71D10) rabbit monoclonal antibody purchased from CST (Cell Signaling Technology). For the measurements, the experimental slides covered with Tks4 fragments were incubated with Myc-CD2AP recombinant protein. The screening experiments and signal visualization were performed by Diagnosticum Zrt using a sandwich-like format in a 300 ul reaction volume in an incubation chamber. First, slides were blocked with SmartBlock solution (CANDOR Bioscience, 113125) for 1 h at 30 °C. Next, the experimental slides were incubated with recombinant Myc-CD2AP protein for 1 h at 30 °C in SmartBlock solution in a final volume of 300 ul. The control slide and the experimental slides were washed four times in 300 ul TBS buffer for 10 min at room temperature then incubated with 1 ug/ml Myc-tag Alexa Fluor 488 antibody in SmartBlock solution in a final volume of 300 ul volume for 1 h at 30 °C. To remove unbound fluorescent antibodies, slides were washed four times with TBS buffer for 10 min at room temperature, and the slides were dried via centrifugation in 50 ml falcon tubes at 1200 rcf for 2 min. Recombinant protein binding was detected by measuring the fluorescence intensity of each peptide spot using a Molecular Devices Axon Genepix 4300A laser scanner. The results were analysed with PepSlide Analyzer Software (Sicasys). The amino acid sequences of the 15-mer TKS4 peptides (with 11 overlapping residues) and the measured fluorescence intensity of the peptide spots on the peptide array are listed in Supplementary Table 3. The relative surface accessibility (rASA) values of each amino acid of the TKS4 fragments were also calculated. For this, the DSSP method (Kabsch and Sander 1983) was applied on the AlphaFold2-derived (Tunyasuvunakool et al. 2021) structure of TKS4 and the absolute surface accessibility values provided by DSSP were divided by the respective maximum surface accessibility values of the amino acids as proposed previously (Tien et al. 2013). Subsequently, the 15mer TKS4 fragments were classified into three accessibility groups based on the number of their residues with rASA values >0.5 to evaluate their position relative to the surface of the folded TKS4 protein and thus the possibility that they can be accessed by CD2AP.

#### 2.9.1. Tissue cDNA array and database analyses

The Colon Cancer Tissue cDNA Array IV panel (HCRT104) was purchased from Origene Technologies (Origene Technologies Inc., Rockville, MD). Each plate contains 48 samples from patients with cancer whose clinical pathological features are freely available at the following link from Origene: https://www.origene.com/catalog/tissues/tissuescan/hcrt104/tissuescan-colon-cancer-cdna-array-iv. The three cecum samples (one from a normal group and two from a tumorous group) were excluded from the analyses, as we were interested in colon tissue only. qRT-PCR was performed with FastStart Green PCR Master Mix, and gene expression levels were normalized to the *GAPDH* and *Pum1* levels (Vermani et al. 2020) based on the method of Hellemans and colleagues (Hellemans et al. 2008). Database analyses were performed using the GENT2 database (Park et al. 2019) and The Cancer Genome Atlas Colon Adenocarcinoma database (TCGA-COAD) using the Xena Platform (Goldman et al. 2020). *CD2AP* and *TKS4 (SH3PXD2B)* mRNA expression level data were collected from tumour or matched normal tissues. From the GENT2 database, datasets including the same nomenclatures (normal, stage I, stage II, stage III and stage IV samples) were exported. Thus, GSE21510 and GSE32323 datasets were omitted from our analyses as these contained A or B sub-stages. The protein interaction network of TKS4 was created using the STRING database (Szklarczyk et al. 2021).

#### 2.9.2. Statistical analyses

Statistically significant differences were calculated using Student’s t-test (p<0.05). Analysis of variance (ANOVA) followed by the Tukey HSD (Honest Significant Difference) test were used for statistical analysis. ROC curve analyses were performed using the Wilson/Brown method. All statistical analyses were performed in GraphPad Prism 8.0.1 (GraphPad Prism, Inc. San Diego, CA).

## 3. Results

### 3.1. CD2AP is a novel TKS4 interacting partner

To shed light on the potential TKS4-CD2AP connection, a high-throughput approach was performed using immunoprecipitation (IP) and mass spectrometry (MS) on several cell lines. The identified TKS4-interacting partners including CD2AP, are shown in Suppl. Table 1. CD2AP protein consistently appeared as a TKS4-associated partner in all analysed cancer cells, i.e., breast (MCF-7), colon (HCT 116, DLD-1), and pancreas (HPAC) cancer types (Figure 1b, Suppl. Table 1.). Furthermore, CD2AP was also present in the precipitate in two non-cancerous cell lines (NIH 3T3, HEK 293)(Suppl. Table 1.). In this study, we focused on the CD2AP-TKS4 interaction; therefore, we sought to further confirm the TKS4-CD2AP binding via IP-western blot (WB) and immunocytochemistry-based microscopy.

The presence of the CD2AP-TKS4 interaction in cancer cells was detected with CD2AP antibody after western blotting of immunoprecipitated TKS4 (Figure 1b) in HCT116 and MCF-7 cells.

To test the localization of CD2AP and TKS4 in HCT116, we have performed single and double staining via immunocytochemistry (ICC) (Figure 1c). The results showed cytoplasmic localization of CD2AP and TKS4. Moreover, the (CD2AP/TKS4) double staining also showed high colocalization events in ICC (Pearson’s coefficient = 0,8172).

To further prove the colocalization of CD2AP and TKS4, proximity ligation assays (PLA) were performed. The PLA also visualized the resulting CD2AP-TKS4 complexes under microscopy in fixed HCT116 colon cancer cells (DuoLink). Micrographs of the assays revealed that CD2AP and TKS4 are in close proximity (at a distance of less than 40 nm) as indicated by the appearance of red fluorescent dots in the cytoplasm and under the cell membrane of the HCT116 cells (Figure 1d).

Next, we decided to map the interaction sites to determine the location of the CD2AP-binding in TKS4 by performing a Pepstar Peptide Microarray analysis. Assuming that one or more of the folded SH3 domains of CD2AP bind proline-rich short linear motifs in TKS4, the binding of full-length CD2AP protein was screened against fragments containing 15mer peptides covering the entire TKS4. To evaluate the accessibility of the peptides within the context of the full-length TKS4 protein, we calculated the relative accessible surface area (rASA) of each amino acid in the TKS4 fragments that showed signs of potential binding. High rASA values indicate that the amino acids are located at the surface of the folded TKS4 (the AlphaFold2-predicted TKS4 structure was used for calculations); thus, they are accessible for binding partners. The results of microarray analysis showed that peptides 188 and 189 of TKS4 showed the strongest interactions with full-length CD2AP and had the highest surface accessibility values (10 of 15 peptides with >50% rASA). Both TKS4 peptides contain a known short linear motif ^755^PVVPPR^760^ that is an optimal binding site for CD2AP SH3 domains (Figure 2). The other TKS4 peptides that bound CD2AP mainly reside within the four SH3 domains of TKS4 with low or medium surface accessibility values (Figure 2), thus these binding events are probably stemming from domain fragmentation and exposure of their sticky hydrophobic cores. Based on our results, we conclude that CD2AP is a novel direct TKS4-interacting partner and that it binds to the ^755^PVVPPR^760^ motif in TKS4 through its SH3 domains.

**Figure 2.**
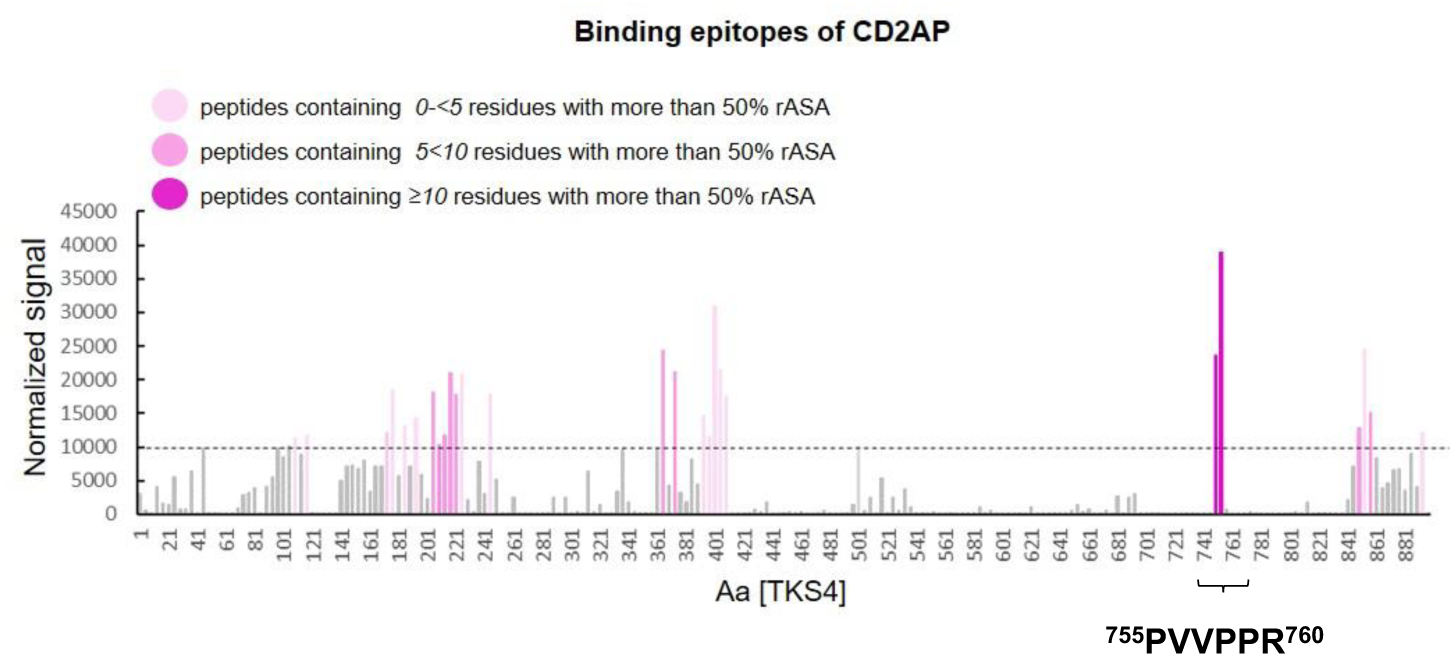
Identification of potential CD2AP binding sites within TKS4. Columns indicate the detected normalized fluorescent signal from the peptide microarray, and the dashed line represents the threshold signal intensity (10000 U). Signals are represented as a mean of three replicates (n=3). The X-axis shows the TKS4 fragments numbered by the position of their first amino acid (Aa) residue in the TKS4 protein sequence. Pink columns represent the normalized signal above the threshold (dashed line). The pink colour shading indicates the degree of the relative surface accessibility (rASA) value (number of residues with >50% rASA) of each 15mer peptide. (*Light pink* columns represent peptides containing *0-<5* residues with more than 50% rASA, *medium pink* columns represent peptides containing *5<10* residues with more than 50% rASA, *dark pink* columns represent peptides containing *≥10* residues with more than 50% rASA.)

### 3.2. Absence of CD2AP and TKS4 promote colon cancer cell migration separately and simultaneously through the partial epithelial-mesenchymal transition (pEMT) process

Although it was already reported that TKS4 regulates cell migration and plays a role in EMT-like process in colon cancer cells (Szeder et al. 2019), and CD2AP has also been connected to cancer development and the inhibition of metastasis in gastric cancer, the underlying mechanisms remained elusive. Therefore, we hypothesized that the TKS4/CD2AP complex cooperatively regulates cytoskeletal rearrangements during cancer progression and participates in EMT process initiation. To investigate the cooperative role of the CD2AP/TKS4 adhesion-related adapter proteins in colon cancer progression, *CD2AP* and *TKS4* were silenced via siRNA treatment either separately or together in HCT116 colon cancer cells, and cell motility assays were performed in the temporarily modified cells. To explore whether CD2AP levels modulate cell migration in functional tests, as was previously reported in the absence of TKS4 protein, wound healing assays were performed. *CD2AP* silencing alone or in combination with *TKS4* silencing enhanced HCT116 cell migration compared with that of the control cells (Figure 3a). Although, no differences have been observed in cell proliferation (Suppl. Fig 1a). We confirmed that CD2AP and TKS4 protein levels were efficiently knocked down after the siRNA treatments (Figure 3b).

**Figure 3.**
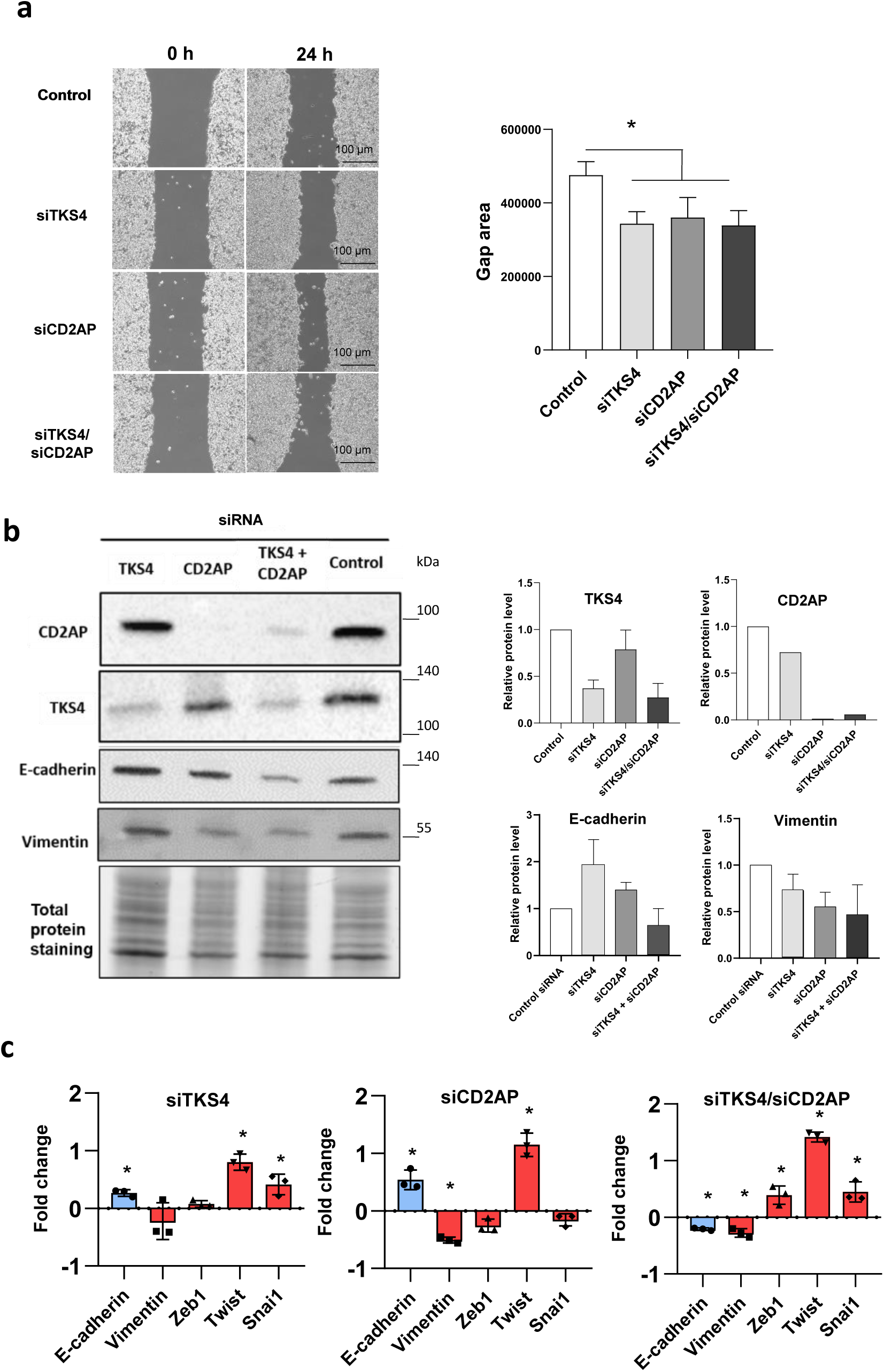
Depletion of *TKS4* and/or *CD2AP* expression in HCT116 colon cells. **(a)** Wound healing assay in *CD2AP*-, *TKS4*-, and *CD2AP/TKS4*-silenced HCT116 cells (upper panel). Between-group comparisons were performed using ANOVA and Tukey’s test, and statistical significance was set at *P<0.05. (n=3) Scale bar=100 µm; Gap area (µm) at 24 hours (lower panel) **(b)** EMT marker protein levels as measured via western blotting (n=3) (E-cadherin and vimentin) and validation of efficient silencing for CD2AP and TKS4. Densitometry analysis for E-cadherin and vimentin. **(c)** qPCR analyses of EMT markers in si*TKS4*-, si*CD2AP*- and si*CD2AP/TKS4*-treated HCT116 cells. Blue bars represent epithelial, red bars represent mesenchymal marker genes. Fold changes show the expressions relative to the mRNA level in negative control siRNA-treated cells (n=3). P values were calculated with Student’s t-test, and statistical significance was set at *P<0.05.

To further examine whether increased cell migration stems from an EMT process, we measured the expression levels of EMT-related genes. In TKS4-silenced cells, epithelial marker E-cadherin expression slightly increased, while mesenchymal marker vimentin was unchanged (Figure 3b, c). The mRNA levels of EMT-related Transcription factors (TF), *Twist* and *Snai1* were also elevated. Thus, moderate expression changes of EMT-related genes were observed. In CD2AP-silenced cells, the RNA and protein levels of the epithelial marker E-cadherin increased (Figure 3b, c), while the mesenchymal marker vimentin decreased. Besides, we measured a reduced level of *Zeb1 m*RNA compared to the control cells, however, *Snai1* level was unchanged and *Twist* transcripts were upregulated (Figure 3c). In short, concurrent expression of both, epithelial-, and mesenchymal-related genes were observed in CD2AP-silenced cells. Simultaneous silencing of TKS4 and CD2AP resulted in decreased E-cadherin and vimentin levels. However, all analysed TFs (*Zeb1, Twist, Snai1*) mRNA levels were upregulated (Figure 3c). To conclude, different co-expression profiles of epithelial and mesenchymal markers were observed depending on which genes were silenced.

To test the hypothesis that CD2AP and TKS4 levels modify cancer cell motility, HCT116 cells were transduced with *CD2AP*- and/or *TKS4-*overexpressing plasmids. The results showed that individual overexpression of *CD2AP* or *TKS4 resulted* in similar level of cell migration compared to control cells. However, the simultaneously enhanced *CD2AP/TKS4* resulted in slower cell migration (Figure 4a). Besides, we measured slightly decreased cell proliferation in TKS4-overexpressing settings (Suppl. Fig. 1b). We confirmed that CD2AP and TKS4 expressions were efficiently enhanced after transfection (Figure 4b). Regarding EMT-related gene expressions, *CD2AP*-, *TKS4*-, and *CD2AP/TKS4*-overexpressing HCT116 cells all have normal E-cadherin and vimentin protein levels (Figure 4b). Additionally, we measured the expression levels of other EMT-related genes via qRT-PCR (Figure 4c). We found that E-cadherin and Vimentin mRNA levels were unchanged in TKS4 and/or CD2AP-overexpressed cells. However, Twist mRNAs were decreased and Snai1 transcripts were elevated in TKS4-overexpressing settings (Figure 4c).

**Figure 4.**
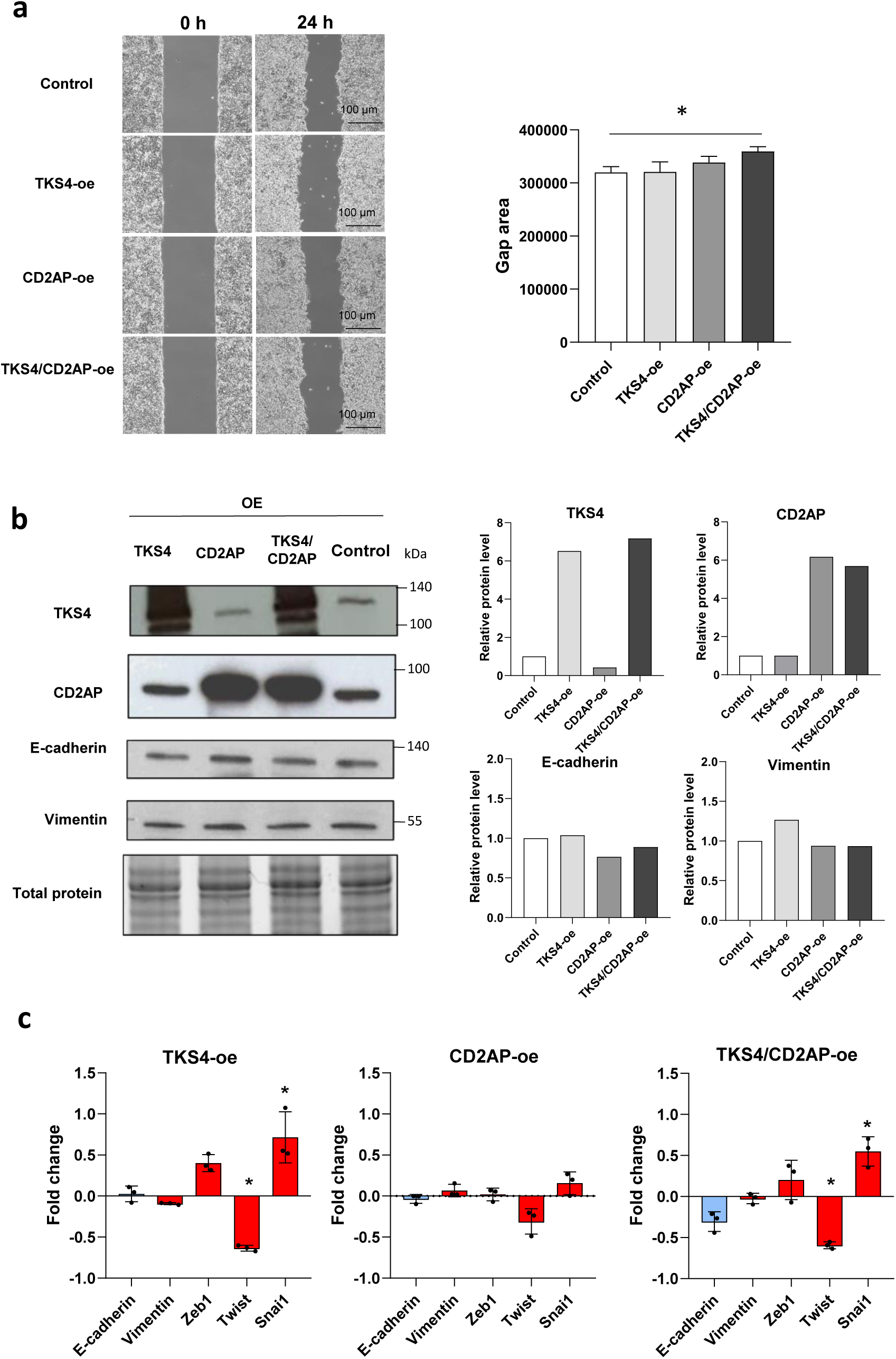
Overexpression of *TKS4* and/or *CD2AP* in HCT116 colon cells **(a)** Wound healing assays with *CD2AP-TKS4-,* and *CD2AP/TKS4-*overexpressing HCT116 cells. (upper panel); Gap area (µm) at 24 hours (lower panel). Between-group comparisons were performed using ANOVA and Tukey’s test, and statistical significance was set at *P<0.05. (n=3) Scale bar=100 µm **(b)** EMT marker protein levels as measured via western blotting. Protein levels of EMT markers (E-cadherin and vimentin). **(c)** qPCR analyses of EMT markers in *TKS4* and/or *CD2AP*-overexpressed HCT116 cells. Blue bars represent epithelial, red bars represent mesenchymal marker genes. Fold changes show the expressions relative to the mRNA level in negative control cells (n=3). P values were calculated with Student’s t-test, and statistical significance was set at *P<0.05.

### 3.4. CD2AP and TKS4 expression levels are useful colon cancer biomarkers

To investigate the expression levels of CD2AP and TKS4 in colon cancer patients, we collected and analysed mRNA data from the GENT2 database. We found lower CD2AP levels and higher TKS4 levels in tumour tissue compared to normal tissue from cancer patients (Figure 5a). To test if further expression pattern changes take place during cancer progression, the GENT2 datasets were grouped to stage I, stage II, stage III and stage IV. While CD2AP levels were equal in all stages, TKS4 expressions were significantly increased in Stage IV compared to Stage II cancer (Figure 5a). We also wanted to validate the results of the database analysis and to further explore the effects of the expression levels on disease progression in human colon cancer patients. Therefore, mRNA expression levels were measured using a qRT-PCR method with a Human Colon Cancer Tissue cDNA Array. Similar to the database results, the tissue cDNA array analysis revealed that the *CD2AP* mRNA level was decreased in cancerous colon tissue compared to normal tissue (Figure 5b). Furthermore, significantly increased *TKS4* expression levels were detected in Stage IV relative to Stage II cancer, and the *TKS4* level increased during disease progression, in agreement with the findings gleaned from the database analyses (Figure 5b). In summary, *CD2AP* level decreases while *TKS4* level increases in colon cancer.

**Figure 5.**
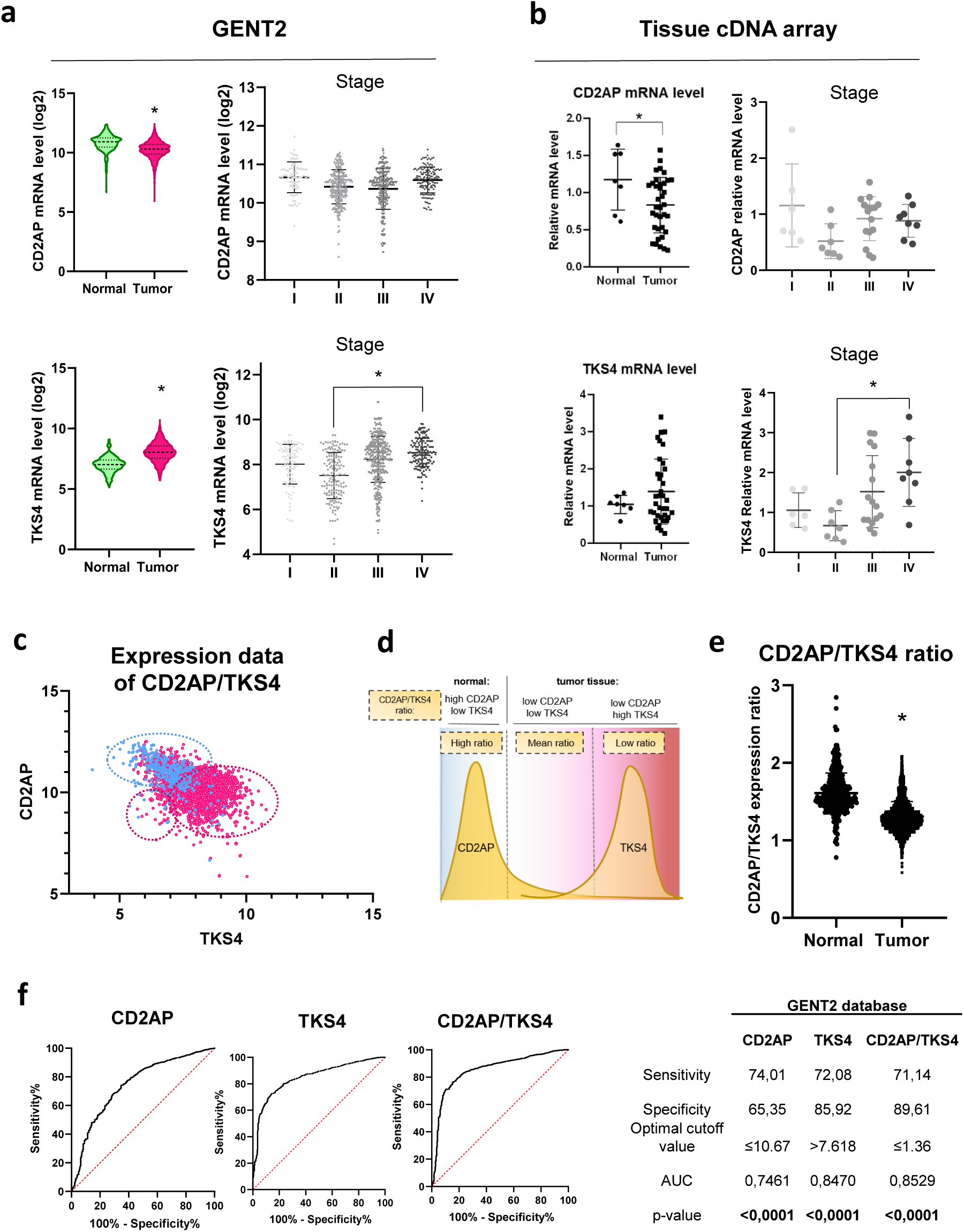
**(a)** *CD2AP* and *TKS4* expression levels in colon cancer patients taken from the GENT2 database (left panel) Normal (n=3553); Tumour (n=3636). *CD2AP* and *TKS4* expression levels in Stage I-IV from GENT2 (right panel) (Stage I (n=77); Stage II (n=422); Stage III (n=304); Stage IV (n=143) **(b)** Colon Cancer Tissue cDNA array results of *CD2AP* and *TKS4* mRNA expression levels in normal (n=7) and tumour samples (n=38) (left panel). *CD2AP* and *TKS4* expression levels in tumour tissue from stage I (n=6), II (n=7), III (n=16) and IV (n=8) cancers (right panel). P values were calculated with Student’s t-test or Tukey’s test (with an ANOVA post-hoc test in the comparison of stages). Statistical significance was set at *P<0.05. **(c)** Merged expression data of *CD2AP* and *TKS4* expression levels in non-tumorous (blue dots) and tumorous colon (pink dots) tissue from the GENT2 database. Transcript levels are shown as gene expression RNA-seq unitlog2(norm_count+1) values. **(d)** CD2AP/TKS4 ratio clusters in normal and tumorous colon tissues. **(e)** The CD2AP/TKS4 gene expression ratio in colon cancer patients from GENT2 database. P values were calculated with Student’s t-test, and statistical significance was set at *P<0.05. **(f)** ROC curve analyses of TKS4, CD2AP and CD2AP/TKS4 ratios from GENT2 data (left panel); Results of ROC curve analyses from GENT2 datasets (right panel).

It was previously reported that the gene expression levels of interacting partners might be correlated (Grigoriev 2001) (Ge et al. 2001), implying that members of the TKS4 interactome might be co-regulated, particularly CD2AP. Therefore, we investigated the relationship between TKS4 and CD2AP expression levels in normal or tumorous tissues. Expression data from GENT2 were plotted to explore whether there is any clustering based on the *TKS4* and *CD2AP* levels. This analysis revealed three clusters defined by different *CD2AP* and *TKS4* levels, namely CD2AP/TKS4 ratios (Figure 5c). Normal samples formed a single “low*TKS4* - high*CD2AP*” cluster (high ratio), while tumour samples formed two distinct clusters: a “low*TKS4* - low*CD2AP*” (mean ratio) and a “high*TKS4* - low*CD2AP*” cluster (low ratio) (Figure 5d). Similar results have been observed investigating data from TCGA-COAD (The Cancer Genome Atlas – Colon Adenocarcinoma database) and cDNA Tissue Array (Suppl. Fig. 2.). Thus, the discrimination of tumorous and normal tissue based on analysis of *CD2AP* or *TKS4* expression alone might result in a false negative detection, while simultaneous analysis of *CD2AP* and *TKS4* expression levels might represents a useful diagnostic marker.

Our results suggest that *CD2AP* and *TKS4* expression levels are associated with colon cancer. As loss of CD2AP or loss of TKS4 alone resulted in different expression patterns of EMT markers *in vitro* (Figure 3), we hypothesized that the *CD2AP/TKS4* expression ratio might have an instructive role in tumour phenotypes. To test this possibility, *CD2AP/TKS4* gene expression ratios were calculated using GENT2 data (Figure 5e). The results showed that the CD2AP/TKS4 gene expression ratio is consistently decreased in tumour tissue (Figure 5e).

To further test the power of the *CD2AP/TKS4* ratio as a biomarker, ROC (Receiver Operating Characteristic) curve analyses were performed based on our database analyses and our colon cancer tissue cDNA array data. ROC analysis is suitable to compare the clinical sensitivity and specificity of particular features, such as *CD2AP* or *TKS4* expression alone as well as the *CD2AP/TKS4* expression ratio in distinguishing normal and tumorous tissue (Figure 5f; Suppl. Fig. 2). The ROC curve analysis of GENT2 data showed that *CD2AP* level is an acceptable marker, while the *TKS4* level, as well as the CD2AP/TKS4 ratio, are excellent markers for distinguishing tumorous and normal phenotypes. For the *CD2AP* and *TKS4* expression levels and the *CD2AP/TKS4* ratio, the area under curve (AUC) values were 0.7461, 0.8470, and 0.8529, respectively (Figure 5f). Of these three features, the *CD2AP/TKS4* ratio was the most specific, with an AUC of nearly 0.9 (Figure 5f, right panel). Similar results have been observed using data from TCGA-COAD and cDNA Tissue Array (Suppl. Fig. 2).

In conclusion, the *CD2AP/TKS4* expression ratio is an outstanding biomarker for the identification of tumorous colon tissue.

## 4. Discussion

Several biomarkers have been developed and used for colon cancer detection; however, due to the high prevalence of this cancer and its poor prognosis, more sensitive and specific biomarkers are needed. Here we show that the previously identified gastric cancer-related CD2AP adaptor protein also has a role in colon cancer development. In addition, based on IP-MS analysis results, we found that CD2AP and the EMT regulator TKS4 form a complex in several different cancer cell types. Besides, we confirmed that CD2AP and TKS4 are in close proximity via Duolink Proximity Ligation Assays.

Via peptide array analysis, we confirmed that CD2AP binds to the ^755^PVVPPR^760^ motif located within the long, disordered linker region between the last two SH3 domains of TKS4 (Figure 1a). The ^755^PVVPPR^760^ sequence in TKS4 represents an optimal recognition motif for the first two SH3 domains of CD2AP. Our results regarding this interaction are reinforced by the results of a previous study that confirmed an interaction between TKS4 and the adaptor protein CIN85 (Bazalii et al. 2015). CIN85 is a paralog of CD2AP, the two proteins show high structural and sequence similarities and their SH3 domains have similar specificities; therefore, we propose that CIN85 likely interacts with TKS4 through binding to the same short linear motif that we identified for CD2AP binding.

We also showed that TKS4 and CD2AP individually and simultaneously modify cell motility and can regulate EMT-related processes in colon cancer. As co-expression of both epithelial and mesenchymal genes were observed in cells lacking TKS4 and/or CD2AP, we suggest that these proteins at normal levels cooperatively inhibit the formation of the pEMT process (Figure 6). Furthermore, different stages of pEMT were observed depending on which genes were silenced. For instance, individual gene silencing resulted in elevated E-cadherin levels, however, some of the mesenchymal-like TFs were upregulated as well. Therefore, we propose that an early pEMT occurs due to a shift in CD2AP/TKS4 ratio stemming from the decreased CD2AP or TKS4 protein levels (Figure 6).

**Figure 6.**
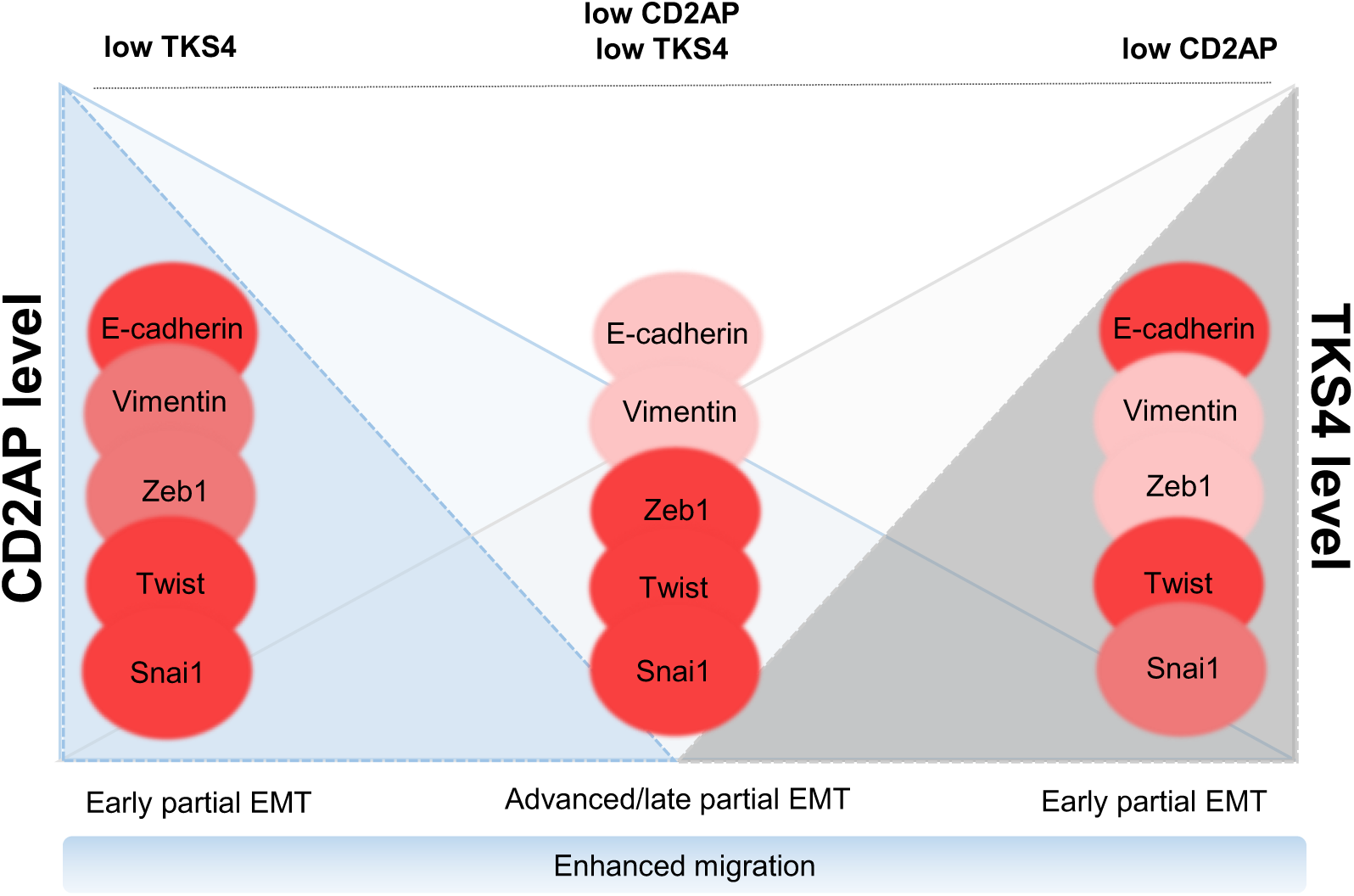
EMT phenotypes, cell migration and expression levels of EMT-related genes grouped according to the levels of the CD2AP and TKS4 proteins based on gene silencing experiments. Shadings of red indicate the expression level changes of the EMT-related gene.

We have manipulated the CD2AP and TKS4 expression levels in the opposite direction and measured the potential mesenchymal to epithelial transition (MET) and the cell motility-related changes in case of overexpression experiments. The CD2AP and the TKS4 overexpression did not result in neither EMT nor MET phenotype. The cell migration changed only in the simultaneous gene-expressing setting and only the Snail TF expression was increased in the TKS4-overexpressed cells compared to the control. These results indicate that epithelial colon carcinoma cells did not gain more epithelial phenotype in case of the overexpression of TKS4 or CD2AP. We also hypothesize that the artificial vector-mediated increased protein levels were valid settings to test the function of the two proteins but could not represent the *in vivo* situation. Supporting this statement on Figure 4, the western blot results showed that the level of TKS4 increased sixfold and showed degradation-indicating bands. In contrast, the results of database analysis showed that in cancer tissue, the median TKS4 increase was less than twice as in the normal sample. Therefore we could not draw a parallel between the shifted CD2AP/TKS4 level seen in cancer patients and the overexpression experiments.

Our results suggest that as adaptor proteins, TKS4 and CD2AP form a complex that organises spatially distinct signal transduction pathways by bringing signalling molecules into close proximity. This type of signal control is not unique, as previous studies have also demonstrated that adaptor proteins (Grb-2, Clb, Crk) can functionally interact with each other to co-organize the members of different signaling pathways (Yablonski 2019)(Buday et al. 1996). As both TKS4 and CD2AP bind to actin cytoskeleton rearranging proteins, we established a model in which TKS4 and CD2AP cooperatively regulate the actin cytoskeleton and actin-based cell motility processes. It was previously confirmed that TKS4 could be membrane-bound (via the PX-domain) and that it binds and co-localizes with cortactin, an actin-binding scaffold protein (Buschman et al. 2009)(Lányi et al. 2011)(Bögel et al. 2012). In addition, CD2AP can also interact with cortactin (Lynch et al. 2003)(Zhao et al. 2013). Moreover, it is already known that cortactin promotes colon cancer cell progression (Ni et al. 2015) and mediates EMT (Ji et al. 2020). Thus, we hypothesize that CD2AP and TKS4 form a stable complex at the cell membrane and that they both conditionally interact with cortactin. This web of interactions could robustly recruit a number of actin-regulating proteins to the membrane to achieve actin remodeling required to create membrane protrusions, which are necessary for the formation of membrane ruffles, lamellipodia, or invadopodia to support metastasis. Although the exact mechanism by which this protein complex regulates EMT is not clear, our current study shows that TKS4 and CD2AP function synergistically inhibit pEMT. Furthermore, as the two proteins play scaffolding roles in actin remodeling, we propose that the TKS4/CD2AP complex might regulate EMT via actin-based cell motility processes.

Previous studies showed that molecular interactions between EMT-related proteins can regulate metastasis. It was proposed that impairing the molecular interactions between adaptor proteins involved in metastasis might be a novel approach for treating aggressive cancer (Feldker et al. 2020). Similarly, our results revealed that the interaction between CD2AP and TKS4 might have a role in tumour development, particularly in regulating a pEMT state in colon cancer cells. Thus, targeting this protein-protein interaction might be a novel approach for inhibiting colon cancer metastasis.

Using cDNA samples from colon tumour samples and RNA-seq data from databases, we found that the *CD2AP* level decreases as the *TKS4* level increases in colon cancer and that further expression level changes can be detected during disease progression. We further found that the CD2AP/TKS4 expression level ratio has an instructive role in tumour phenotypes. We would like to highlight that one situation among our expression level changing functional studies might mimicked the *in vivo* observed situation: when CD2AP was downregulated and the level of TKS4 was unchanged. We found that silencing CD2AP led to enhanced cancer cell migration and concurrent expression of epithelial and mesenchymal markers accompanied with high Twist transcription factor expression (Figure 3). In the case of the CD2AP/TKS4 ratio was artificially shifted the phenotype of the HCT116 cells started to change in the mesenchymal direction. We hypothesize that disturbed EMT signaling regulation induced by the decreased CD2AP/TKS4 expression ratio in colon cancer cells might enhance their migratory capacity and oncogenicity.

Our ROC curve analyses suggest that the CD2AP/TKS4 expression ratio is an outstanding biomarker for the identification of tumorous colon tissue. Further exploration of the molecular interaction between CD2AP and TKS4, their cooperative roles in EMT, as well as the implications of the decreased CD2AP/TKS4 expression ratio in tumours might reveal additional details that could be used to develop novel approaches for detecting colon cancer. From a broader perspective, our study highlights the possibility that gene expression ratios might be more reliable biomarkers than the expression levels of single genes for the diagnosis of early-stage colon cancer and perhaps additional cancer types, such as breast or pancreatic adenocarcinoma.

## Conclusions

Our study revealed that CD2AP is a novel TKS4-interacting protein and together they form a scaffold complex. They function synergistically to inhibit partial EMT as they potentially regulate EMT-related signaling in HCT116 cells. Furthermore, our results highlight that the relative expression levels of multiple genes – as in the case of CD2AP and TKS4 - might serve as more reliable biomarkers than the expression levels of single genes for the diagnosis of early-stage colon cancer. At last, as the interaction between CD2AP and TKS4 might play a role in tumour development, targeting this protein-protein interaction might be a novel approach for inhibiting colon cancer metastasis.

## Additional files

## Ethical approval

Not applicable.

## Author contributions

AK initiated the project, designed the study, carried out the experiments, analysed the data, supervised students, and wrote the manuscript. VV participated in the coordination, conceptualization, and implementation of the study, helped in data analysis as well as writing and editing of the manuscript. LL contributed to the experimental work, especially the DuoLink assay, data interpretation, and manuscript preparation. TT and JN helped in cell-based experiments and in data visualization. ÁT contributed to the analysis steps and supported manuscript preparation. LL contributed to the western blot experiments and data interpretation. RP contributed to data interpretation and visualization, as well as editing the manuscript. LB supervised the project and obtained the funding. All authors read and approved the final manuscript.

## Acknowledgments

We thank Zita Haidar for technical assistance and Éva Csősz for valuable help with the mass spectrometry experiments. The authors thank Rita Lózsa, Dávid Szüts, Gyöngyi Bézsenyi and László Homolya for the technical help and the tasks related to cell lines. We also thank János Fekete for his help with bioinformatic analyses. We are especially grateful to Balázs Győrffy for his useful comments about the manuscript.

## Data Availability Statement

The data and materials supporting the conclusions of this study are included within the article and its additional files.

## Additional files

**Supplementary Figure 1.** Cell proliferation assays.

**Supplementary Figure 2.** (A) The CD2AP/TKS4 gene expression ratio in colon cancer patients from the TCGA-COAD database and (B) based on the Colon Cancer Tissue cDNA array results. P values were calculated with Student’s t-test, and statistical significance was set at *P<0.05. (C) ROC curve analyses of TKS4, CD2AP and CD2AP/TKS4 ratios from the TCGA-COAD database. (D) ROC curve analyses of TKS4, CD2AP, and TKS4/CD2AP ratios to discriminate normal and tumour tissue from the Colon Cancer Tissue cDNA array. (E) Results of ROC curve analyses from TCGA or Tissue cDNA array datasets.

**Supplementary Table S1.** List of TKS4 interacting partners

**Supplementary Table S2.** Oligos used in the study

**Supplementary Table S3.** TKS4 and CD2AP peptide array. PX and SH3 domains are shown with bold letters and signals up to 10000 UA are grey.

