## Supplementary material for "Exploring the Interdependence of TKS4 and CD2AP: Implications for EMT Process and Early Detection in Colon Cancer": Supplementary Methods.docx

**Cell proliferation assays**

Cells viability were measured via MTT assay using Cell Proliferation Kit I (MTT) - Cat. No. 11 465 007 001- Roche. 5 x 10^3^ HCT116 cells were seeded in a 96-well plate in a final volume of 100 μl culture medium per well. The same day, cells were transfected with siRNAs or plasmids then cell numbers were counted after 24 and 48 hours. After the appropriate incubation period 10 ul MTT labelling reagent was added to the wells and we incubated for 4 hours at 37°C 5% CO2. Then 100 ul Solubilization reagent was added to each well and incubated overnight 37°C 5% CO2. Next day, we measured the absorbance with an ELISA plate reader at 600 nm and a reference absorbance at 700 nm for background measurements.
