## Supplementary figures and images for "Exploring the Interdependence of TKS4 and CD2AP: Implications for EMT Process and Early Detection in Colon Cancer"

### Suppl. Fig. 2..JPG

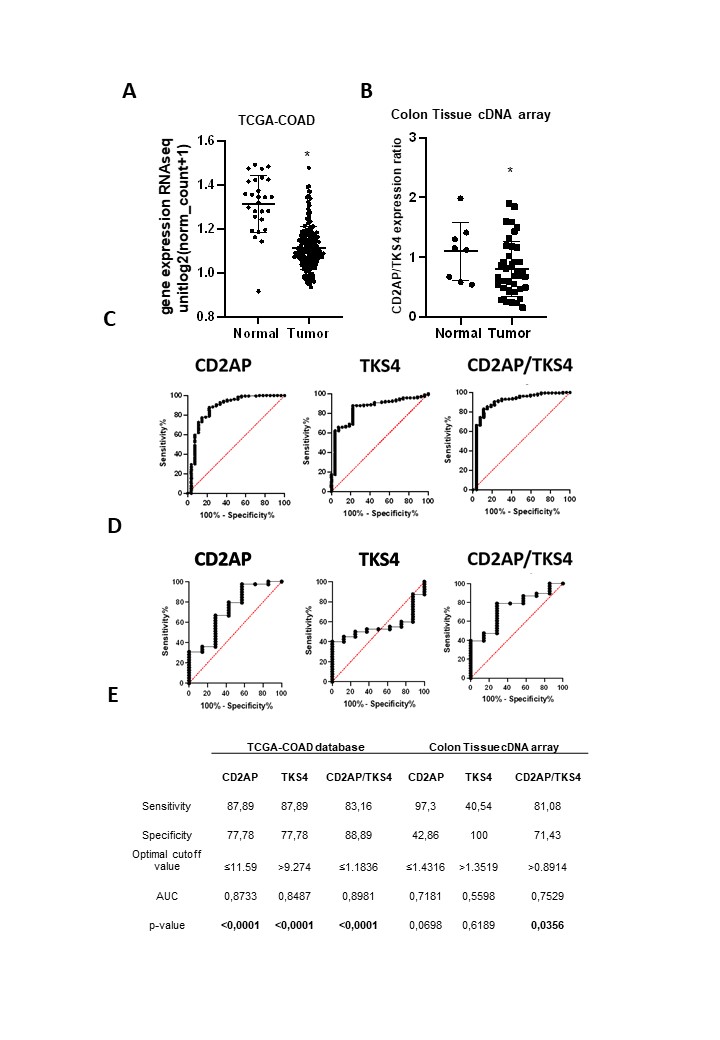
